# Aggregation of Cohorts for Histopathological Diagnosis with Deep Morphological Analysis

**DOI:** 10.1101/2020.10.13.337014

**Authors:** Jeonghyuk Park, Yul Ri Chung, Seo Taek Kong, Yeong Won Kim, Hyunho Park, Kyungdoc Kim, Dong-Il Kim, Kyu-Hwan Jung

## Abstract

There have been substantial efforts in using deep learning (DL) to diagnose cancer from digital images of pathology slides. Existing algorithms typically operate by training deep neural networks (DNNs) either specialized in specific cohorts or an aggregate of all cohorts when there are only a few images available for the target cohort. A trade-off between decreasing the number of models and their cancer detection performance was evident in our experiments with The Cancer Genomic Atlas (TCGA) dataset, with the former approach achieving higher performance at the cost of having to acquire large datasets from the cohort of interest. Constructing annotated datasets for individual cohorts is extremely time-consuming, with the acquisition cost of such datasets growing linearly with the number of cohorts. Another issue associated with developing cohort-specific models is the difficulty of maintenance: all cohort-specific models may need to be adjusted when a new DL algorithm is to be used, where training even a single model may require a non-negligible amount of computation, or when more data is added to some cohorts. In resolving the sub-optimal behavior of a universal cancer detection model trained on an aggregate of cohorts, we investigated how cohorts can be grouped to augment a dataset without increasing the number of models linearly with the number of cohorts. This study introduces several metrics which measure the morphological similarities between cohort pairs and demonstrates how the metrics can be used to control the trade-off between performance and the number of models.

## Introduction

Pathologists diagnose cancer from hematoxylin and eosin (H&E) stained slides after comprehensively analyzing the histological features in a given slide^1^. Likewise, data-driven algorithms including deep learning (DL) have been devised to detect cancer using H&E morphology^2^. While a larger dataset size used to train DL algorithms directly translates to better performance, previous applications to cancer detection train models on cancer-specific datasets such as breast cancer^2–5^, skin cancer^6–8^, lung cancer^9^, bladder cancer^10^, prostate cancer^8^ and lymph node metastases^8, 11, 12^, with restricted capacity from limited data. Another approach is to develop a universal model agnostic to cohort type with the hope that increasing the dataset size outweighs the drawbacks brought by introducing irrelevant information or features^13^. Our experiments, however, demonstrate that training a universal model to detect cancer from H&E images collected from various cohorts results in worse performance than that achieved by ensembling cancer-specific models. This prompts the following question: when and how can datasets be aggregated to better detect cancer from fewer dataset configurations?

At first sight, it may be tempting to conclude that training models on cohorts with similar histologies would enhance their performance. However, dataset aggregation based on subjective evaluation of morphological similarity can result in inclusion of irrelevant information that outweighs the benefits brought by diversity. It would also be desirable if cohorts could be grouped objectively by similarity without pathologists’ expertise. In this study, we show how cohort aggregation can be done appropriately without any domain knowledge by introducing several metrics inspired by domain adaptation^14^ (see Materials and methods), and that the performance of DNNs can be improved with a reduced number of models.

We explored the effect of combining datasets belonging to different cohorts in developing a cancer diagnostic model from H&E images. Our study was conducted using The Cancer Genome Atlas (TCGA) dataset comprised of 37 cohorts across 33 cancer types^15^. This dataset has been used in various studies for stain color handling^16^, microsatellite instability prediction^17^, tissue classification^18^ and gene mutation prediction^9^. We included 12 TCGA cohorts which contain at least 36 normal slides: (KIRC, Kidney renal clear cell carcinoma; LIHC, Liver hepatocellular carcinoma; THCA, Thyroid carcinoma; OV, Ovarian serous cystadenocarcinoma; LUAD, Lung adenocarcinoma; LUSC, Lung squamous cell carcinoma; BLCA, Bladder Urothelial Carcinoma; UCEC, Uterine Corpus Endometrial Carcinoma; BRCA, Breast invasive carcinoma; PRAD, Prostate adenocarcinoma; COAD, Colon adenocarcinoma; STAD, Stomach adenocarcinoma) as summarized in Table 1. Note that some cohorts in this dataset such as COAD and STAD are known to share similar morphologies as illustrated in Figure 1. After validating the diagnostic performances of cohort-specific models and comparing their performance with that of a universal model trained on all 12 cohorts, we show how a careful aggregation guided by our metrics can be used to reduce the number of models while retaining the high performance achieved by the specialized models. In one aspect, our work is an extension of that conducted by Kather *et al*.^17^ who trained a microsatellite instability (MSI) detection model on gastric formalin-fixed paraffin-embedded (FFPE) slides and validated the model on colorectal FFPE slides, demonstrating the applicability of DL-based classification on morphologically similar cohorts. In contrast to their experiments, we show that a direct usage of both source and target cohorts guided by carefully designed metrics can be used to automate the cohort aggregation process.

**Table 1.** The Cancer Genome Atlas (TCGA) cohorts and the number of frozen slide images used in the study. Cohort abbreviations are as follows: KIRC, Kidney renal clear cell carcinoma; LIHC, Liver hepatocellular carcinoma; THCA, Thyroid carcinoma; OV, Ovarian serous cystadenocarcinoma; LUAD, Lung adenocarcinoma; LUSC, Lung squamous cell carcinoma; BLCA, Bladder Urothelial Carcinoma; UCEC, Uterine Corpus Endometrial Carcinoma; BRCA, Breast invasive carcinoma; PRAD, Prostate adenocarcinoma; COAD, Colon adenocarcinoma; STAD, Stomach adenocarcinoma

| Cohort | Number of positive slides | Number of negative slides |
| --- | --- | --- |
| KIRC | 1089 | 564 |
| LIHC | 399 | 89 |
| THCA | 534 | 97 |
| OV | 1175 | 163 |
| LUAD | 819 | 244 |
| LUSC | 753 | 347 |
| BLCA | 431 | 37 |
| UCEC | 750 | 54 |
| BRCA | 1572 | 399 |
| PRAD | 604 | 118 |
| COAD | 871 | 109 |
| STAD | 632 | 123 |

**Figure 1.**
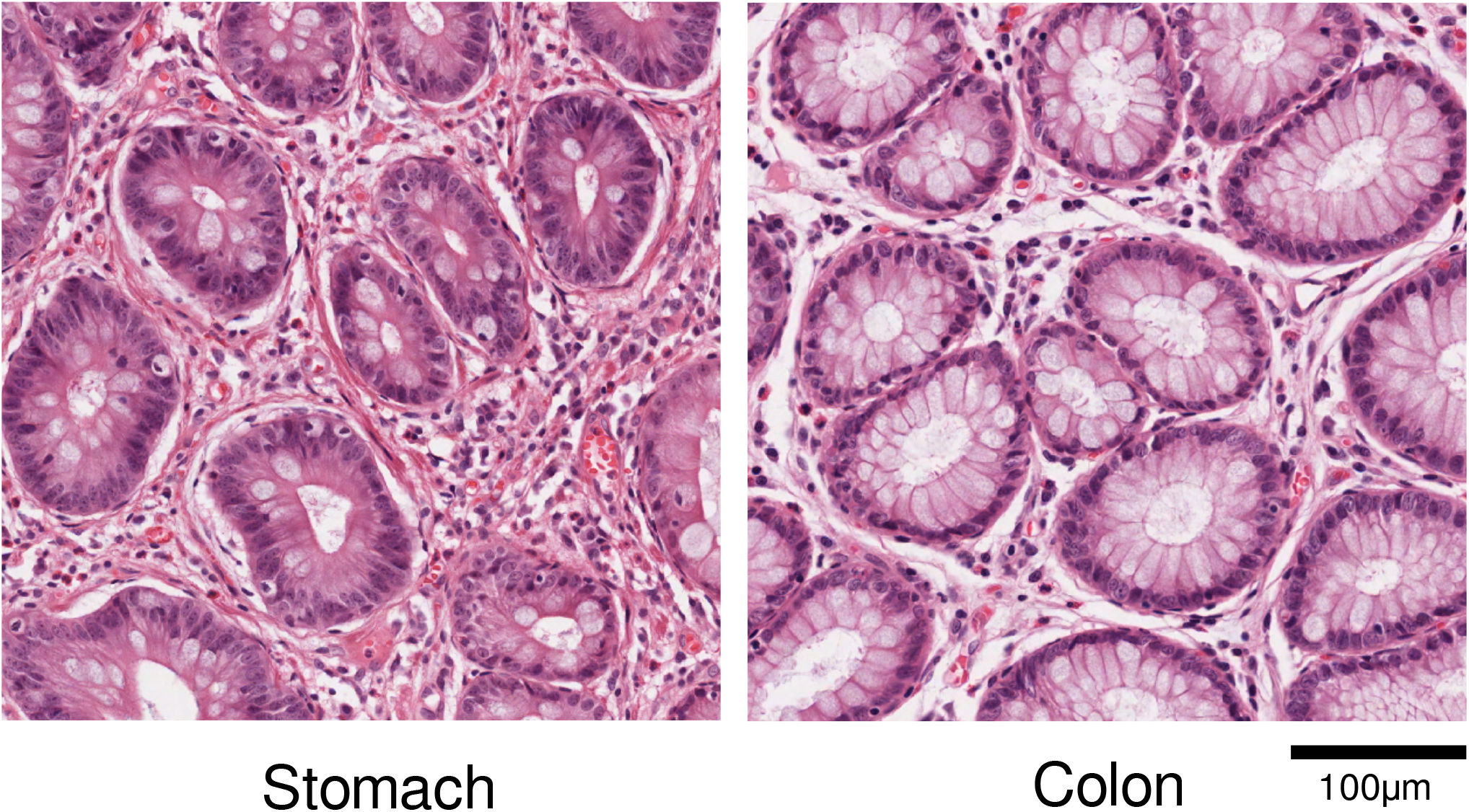
Examples of H&E patch from gastric and colonic tissues. On the left are gastric glands with intestinal metaplasia, and on the right are colonic glands. The two tissues share similar histological features.

## Results

We conducted several experiments using the dataset and model setups (Figure 2 and Figure 3) to test if prior histology knowledge of morphological similarities among different cancer types could be used to train DL models efficiently. Our first experiment demonstrates the sub-optimality of using a single universal model trained on all H&E images regardless of their cohorts of origin. Three DL models were then trained to discriminate a slide’s cohort of origin based on positive (cancer present), negative (cancer absent), or either type of slides, where intuitively, they would have difficulty distinguishing cohorts with similar morphological structures. This idea stems from the 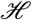-divergence introduced in the context of domain adaptation^14^, where the difficulty of discriminating between two domains (i.e. cohorts) can be used to understand the feasibility of training on a source cohort for the task on a target cohort. The models obtained from these experiments were then used to guide the aggregation of cohorts. Our aggregation method allows controlling the trade-off between performance and number of models, with models trained on cohort groups generally achieving higher performance as the number of models increase (i.e. with higher specialization).

**Figure 2.**
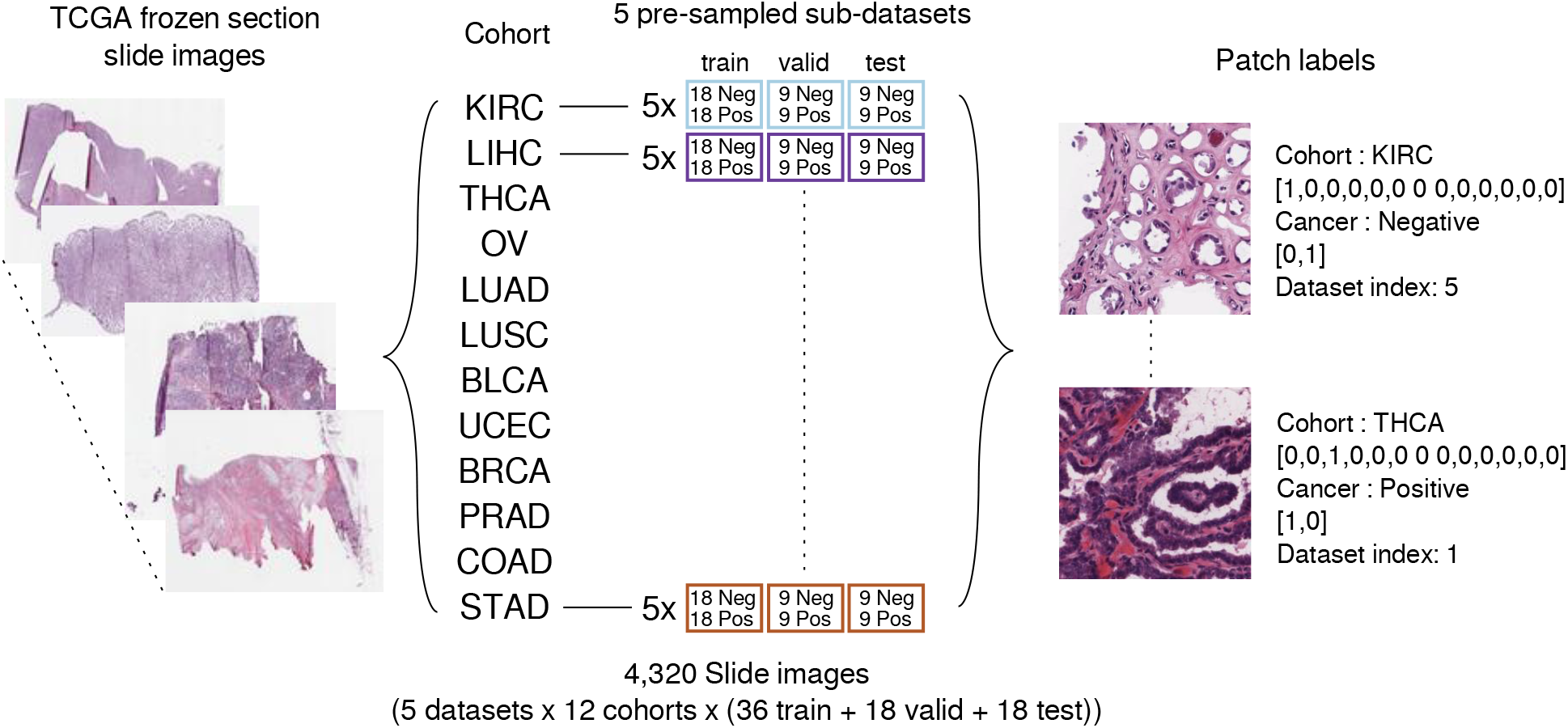
Dataset preparation: a total of 4,320 frozen slide images were used for train/validation/test. Five sub-datasets were constructed for each cohort, with each sub-dataset composed of randomly sampled slides numbering (cancer detection and general cohort discrimination) 18 positive and negative slides or (positive/negative cohort discrimination) 36 positive or negative slides, with replacement between sub-datasets. Each patch was annotated indicating its cohort, presence of cancer, and sub-dataset index.

**Figure 3.**
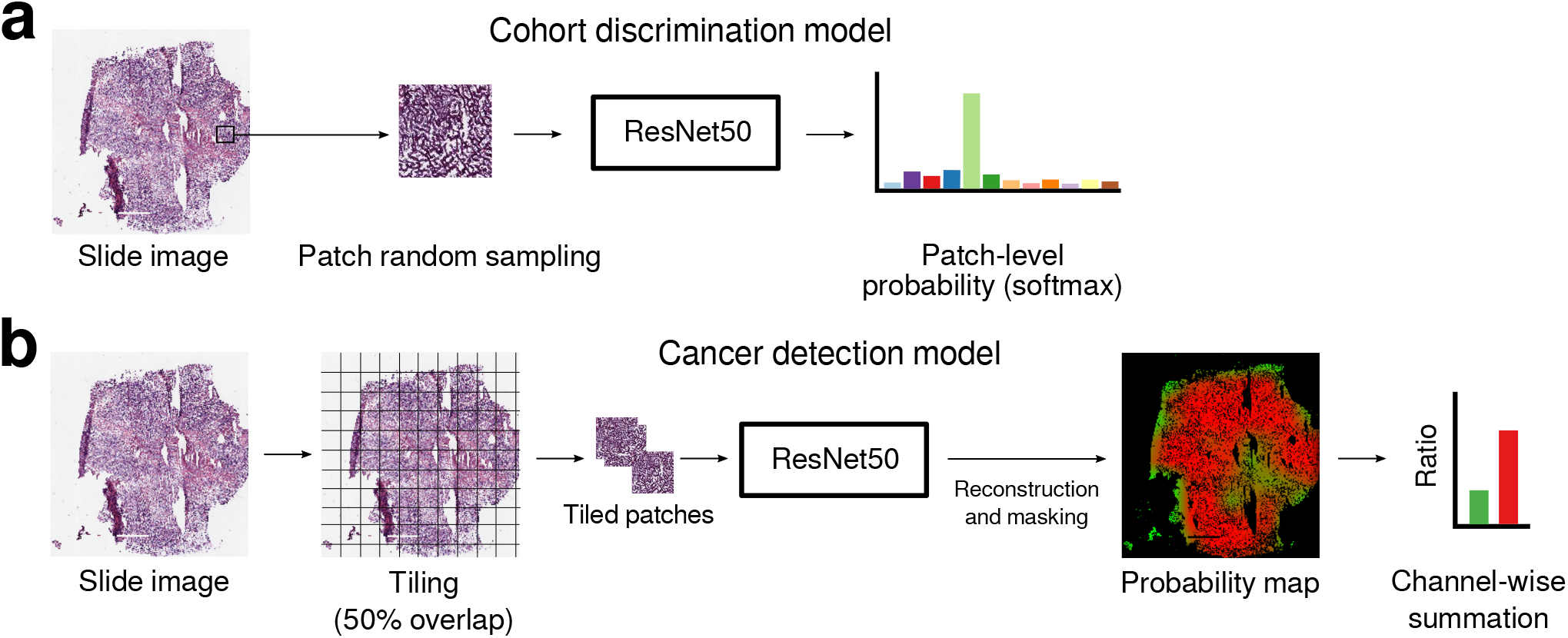
Input/output schematic of deep learning models. Cohort discrimination model (a) predicts which cohort a patch is drawn from. Cancer detection model (b) makes slide-level predictions determining whether the slide contains cancer or not.

### Specialized and universal cancer detection models and morphological similarities among cohorts

To compare the performances of cohort-specific and universal cancer detection models, we first trained 12 cohort-specific models to diagnose cancer from H&E images in designated cohorts. A universal model was then trained on an aggregate of all 12 cohorts, and its area under the receiver operating characteristic (AUROC) scores were obtained when tested on each of the 12 cohorts. Indeed the average AUROC of cohort-specific models (the average AUROC 0.9687) far exceeds that of the universal model (the average AUROC 0.8570), demonstrating how merging cohorts with distinct morphologies yields highly sub-optimal performance. Furthermore, we tested all cohort-specific models on all cohorts regardless of their source cohort (training set) to obtain a 12 × 12 similarity matrix (Figure 4a). This similarity matrix gives a proximity measure between cohort pairs, which can in turn be used to group cohorts using hierarchical clustering analysis (HCA).

**Figure 4.**
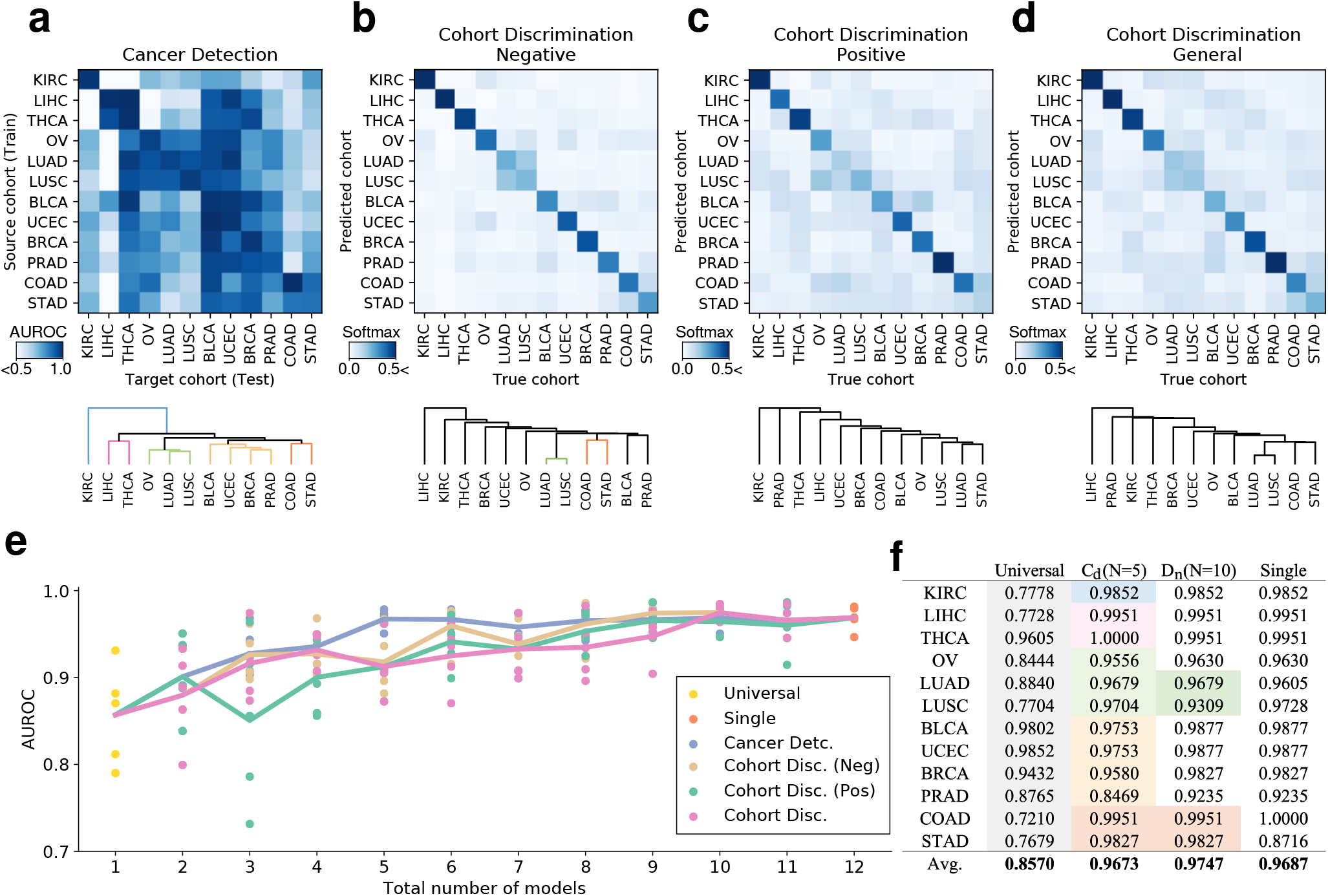
Similarity matrices according to four deep learning models and corresponding hierarchical clustering analysis (HCA) results. Single-cohort cancer detection models *C_d_* (a) with rows indicating train (source) cohorts and columns indicating test (target) cohorts. Negative cohort discrimination model *D_n_* (b) with rows indicating cohort of origin and columns indicating the model’s predictions. Positive *D_p_* (c) and general *D_g_* (d) cohort discrimination models. A few lines in the HCA grouping are colored either by cohort (KIRC and LIHC) or organ (lung: LUAD and LUSC, GI tract: STAD and COAD) for tracking. (e) Performances of DNNs trained on all possible aggregations of cohorts according to HCA groupings obtained from different similarity matrices (a-d). All AUROC scores are shown with lines connecting the means. (f) AUROC scores of universal, N = 5 models with cohorts grouped according to single cancer detection models, N = 10 models grouped according to negative discriminative model, and N = 12 models specialized for each cohort. Cohorts are colored consistently with their respective lines in HCA.

The diagonal entries in the similarity matrix show that cohort-specific models perform best when the target cohort matches the source cohort as expected. The low AUROCs in off-diagonal entries show the degree of incompetence of cohort-specific models when tested on other cohorts. Cohort groupings obtained from HCA show that KIRC is unique, standing out from the other cohorts of interest. LUSC and LUAD were determined most similar, implying high morphological similarity between the two.

### Cohort discrimination models and morphological similarities among cohorts

Next, we measured the morphological similarities among cohorts using cohort discrimination models. Three models were trained to identify which cohort the H&E image belongs to, when trained on different image types: (1) a positive discrimination model *D_p_* trained on positive (cancer present) images, (2) negative discrimination model *D_n_* trained on negative (cancer absent) images, and (3) general discrimination model *D_g_* trained on both positive and negative images. Before presenting our main results, we describe the intuition connecting the cohort discrimination task in determining which cohorts are similar, which in turn is used to aggregate cohorts and control the trade-off between performance and number of models. Pathologists agree that some organs (e.g. stomach and colon, Figure 1) share similar histological structures that may not be easily distinguished to the untrained eye. If this is an inherent difficulty that cannot be overcome by any expressive model, DNNs would also struggle in discriminating such images, and adding the extra images from another cohort would be similar to simply increasing the number of training samples in the original cohort of interest without any expense of adding irrelevant information. To quantify the similarity among cohorts, positive-, negative-, and general cohort discrimination models were trained to identify which cohort positive, negative, or either H&E patches were retrieved from, respectively.

These cohort discrimination models’ one-versus-all classification performances are displayed in Figure 4b-d along with their HCA groupings. We had expected the negative cohort discrimination model *D_n_* to especially struggle with distinguishing cohorts collected from the same organ (e.g. lung cohorts LUSC and LUAD), as the negative H&E images do not have cancer-related patterns which is useful in differentiating the organ of origin. As expected, COAD-STAD and LUSC-LUAD which are biologically the similar or originate from the same organs were grouped together when cancer was absent (Figure 4b), showing that gastrointestinal tract cohorts and the two lung cohorts were similar in morphology. The former pair was also grouped together by the positive discriminative model (Figure 4c), implying their morphological similarity even when cancer is present. This observation is in line with clinical knowledge that pulmonary adenocarcinomas can be difficult to distinguish from pulmonary squamous cell carcinomas when it histologically shows poor differentiation. The liver and kidney renal clear cell carcinoma have unique tissue structures (Figure 5), and we had expected the cohort discrimination models to distinguish these cohorts with high accuracy, and the resulting HCA grouping to isolate these cohorts. The negative discriminative model isolated LIHC (Figure 4b), and similarly, the positive discriminated isolated KIRC (Figure 4c), indicating H&E images from LIHC and KIRC with and without cancer, respectively, have substantially different morphologies from other cohorts.

**Figure 5.**
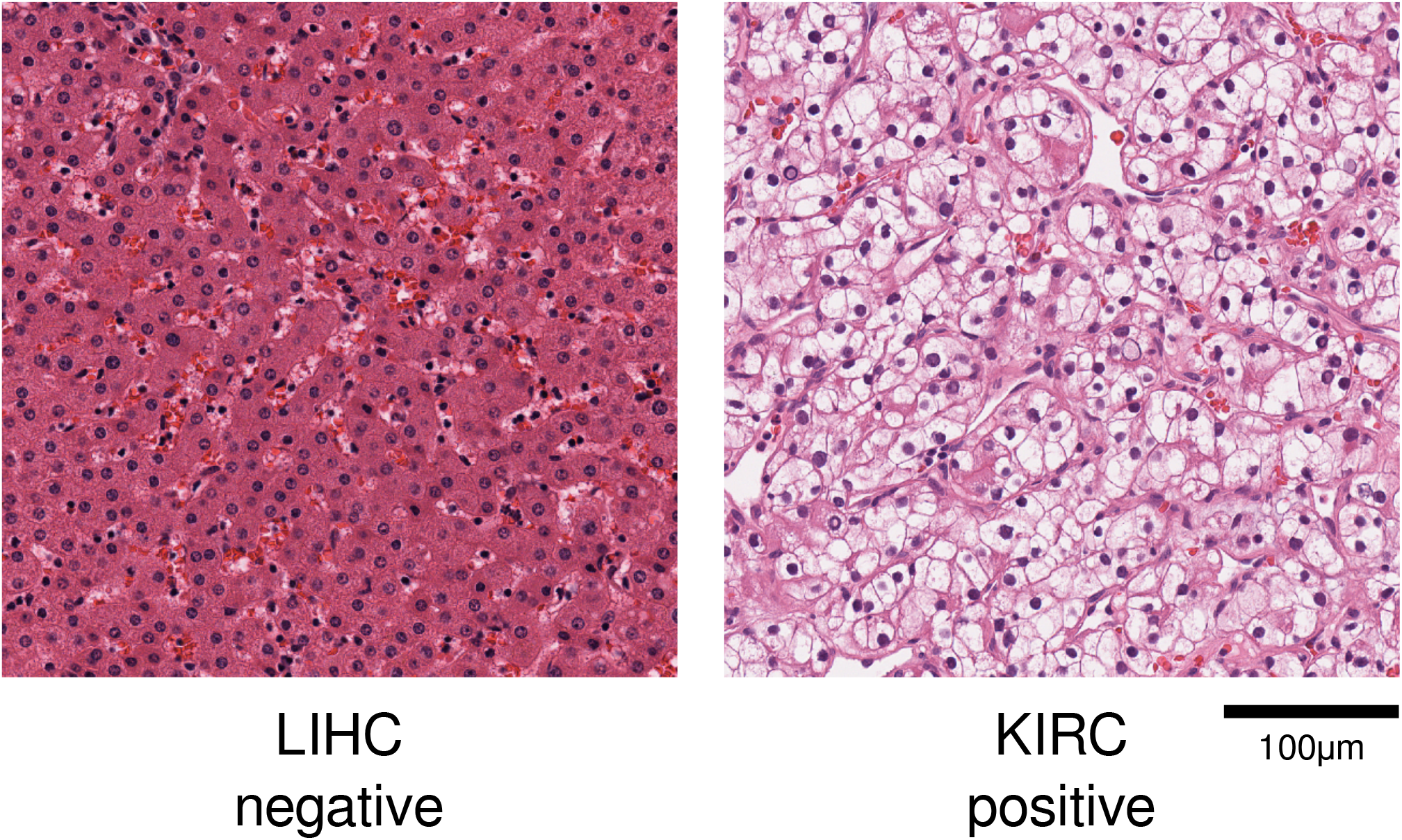
Representative images of the morphological features unique to LIHC negative and KIRC positive slides are shown. LIHC (negative) shows cords of reddish polygonal hepatocytes, and KIRC (positive) shows clear cell clusters with distinct cellular borders.

### Qualitative analysis of morphological similarity among cohorts

To qualitatively analyze the similarities among cohorts, we visualized the features extracted by the universal model and general cohort discrimination model using Uniform Manifold Approximation and Projection (UMAP) (Figure 6; Materials and methods)^19^ Both UMAP visualizations projected features from STAD and COAD close to each other; however, there was still a clear distinction between the two. This demonstrates how STAD and COAD are similar in morphology, but have unique traits not shared between the two. THCA and BRCA are on the other end of the spectrum, with the pair being distinguished from others, but their features overlapping substantially. Both the universal cancer detection and general cohort discrimination models isolated KIRC, indicating that KIRC shares the least amount of morphological properties with other cohorts. Surprisingly, while LIHC has a unique hepatic tissue structure absent in all other cohorts considered, it is only isolated when features were extracted using the cohort discrimination model and is rather uniformly distributed when extracted by the universal model. In general, the clusters formed by the universal model’s features was consistent with that obtained from the cohort discrimination model explicitly trained to discern cohorts, even though the universal model had no prior knowledge that the aforementioned cohorts have unique morphological features. The clustering derived from the universal model’s features in the UMAP plot and cohort-specific models’ HCA grouping shared a similar pattern, yielding consistently isolated and neighboring cohorts. This confirms the similarity/dissimilarity among cohorts via independent dimension reduction techniques.

**Figure 6.**
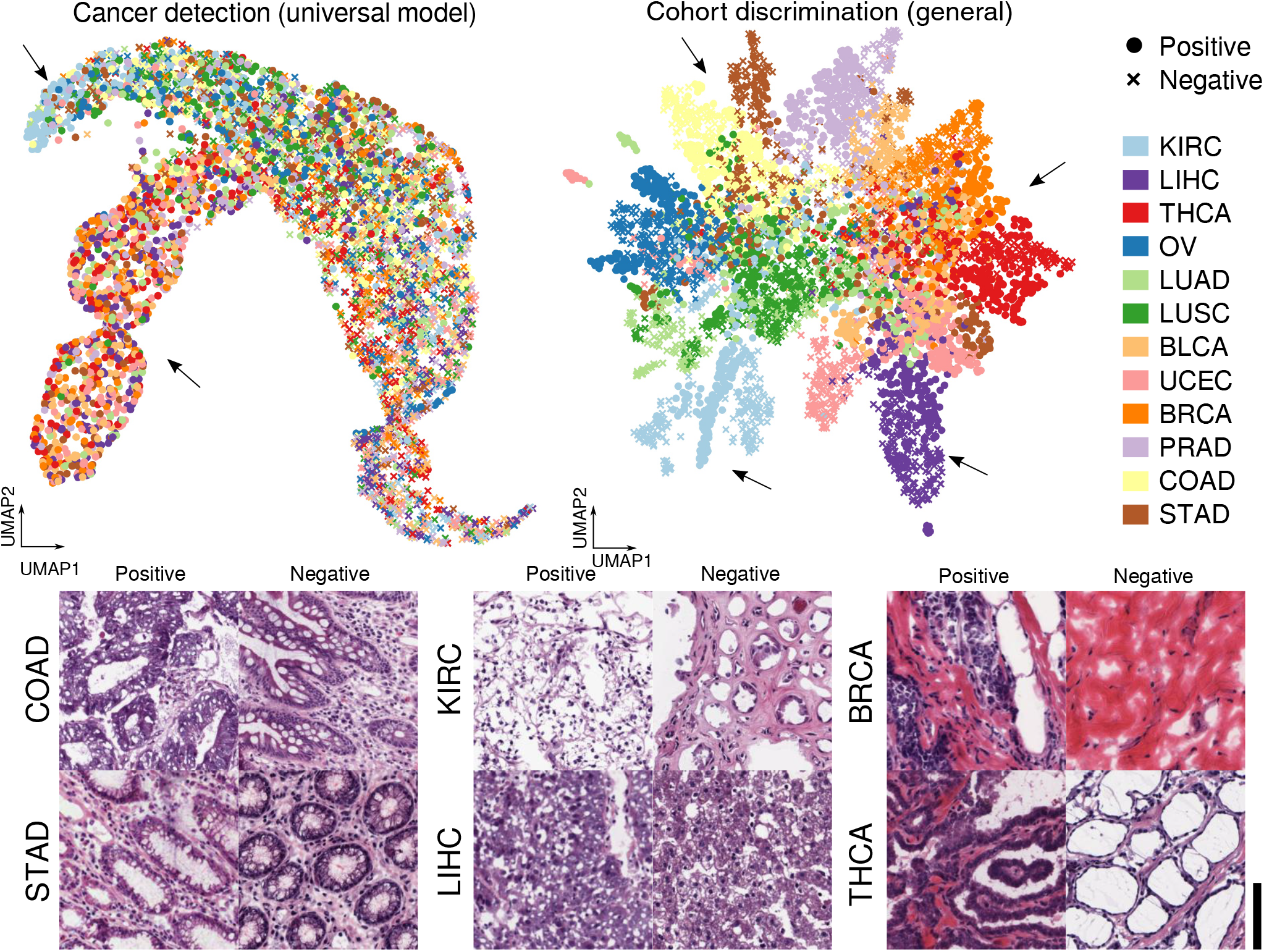
Uniform Manifold Approximation and Projection (UMAP) visualizations for features extracted by the universal cancer detection model (N = 1) and general cohort discrimination model. Identical patches were used for both visualizations. Cohort groups that were clearly isolated from others are indicated with arrows with their corresponding patches shown below: KIRC, LIHC, THCA-BRCA, and STAD-COAD. COAD and STAD have tall tubular glandular structures; KIRC and LIHC have unique cellular and tissue features distinct from other cancers. THCA shows papillary cores with nuclear crowding in a positive patch and a uniform follicular sheet structure in a negative patch. Scale bar: 100*μ*m

### Aggregating cohorts while retaining performance

Using the four aforementioned models and their corresponding measures of cohort similarity, we trained and validated the performance of DNNs trained on cohort groups obtained from the four respective groupings. As shown in Figure 4e, the average AUROC of DNNs generally increased as the models specialized in specific cohorts. When cohorts were grouped according to our measures of similarity, we were able to achieve almost the same performance as that achieved by 12 specialized models using only 5 models. Furthermore, the performance of 10 models trained on an aggregate of the 12 cohorts carefully combined according to the negative and general cohort discrimination models slightly outperformed the specialized models. This demonstrates the need to handle dataset configurations more carefully when applying DL to cancer detection, especially to avoid constructing conclusions based on sub-optimal models.

Instead of exhaustively listing which cohorts were grouped together to train cohort-group models, we list the groupings which resulted in highest performance: groupings obtained by *C_d_* with *N* = 5 models and *D_n_* with *N* = 10. (*C_d_*) {*KIRC*}, {LIHC, THCA}, {OV, LUAD, LUSC}, {BLCA, UCEC, BRCA, PRAD}, {COAD, STAD}, (*D_n_*) {KIRC}, {LIHC}, {THCA}, {OV}, {LUAD, LUSC}, {BLCA}, {UCEC}, {BRCA}, {PRAD}, {COAD, STAD}.

## Discussion

Constructing datasets composed of multiple cohorts with similar morphologies can be useful for the following reasons. First, utilization of a vast amount of publicly available data can increase a DNN’s performance trained using DL algorithms. Second, appropriate aggregating of cohorts can reduce the number of diagnostic models required to diagnose a given histologic slide image: a physican does not need to synchronize the algorithms across cohorts whenever new data is added to the training set. Lastly, it can be used to develop an algorithm for rare cancers for which there does not exist a large number of histologic images, using datasets from more well-known cancer types that bear histologic resemblance.

Previous DL algorithms in digital pathology often used arbitrary combinations of cohort, depending on their availability. Existing histological diagnostic algorithms are either organ-specific or organ-agnostic modules developed without consideration of morphological similarities or dissimilarities among cohorts. To the best of our knowledge, this is the first study to systematically analyze cohort morphologies by cancer type using modern data-driven methods. We trained DNNs on 4 different tasks to express the similarities among cohorts as features extracted from the trained models, which in turn was used to produce hierarchical clusters. The obtained clusters were used to effectively group cohorts such that models trained on these groupings perform as well as or better than both the universal model and the most specialized, single-cohort models.

We address some possible extensions for future work. Our study considered aggregating datasets to improve the performance of cancer detection of a universal and single-cohort models achieved relatively low performances in Figure 4f. A natural extension of the current study would be to see if the proposed metrics still prove useful in detecting cancer using other cohorts of interest. Another course for future work would be applying our framework to other datasets. Concurrent with our work, Hosseini *et al*^20^ published an annotated pathological image database with hierarchical ordering and it would be interesting to see if the cohorts’ properties remain unchanged when our framework is applied to their dataset. Recall that similarity among cancer tissues’ features affects the cancer detection models more than normal organ features: the grouped cohort models’ performances showed a higher correlation with the positive discrimination model’s metric than the negative model’s. This suggests that in order to develop accurate cancer detection models, it is crucial to allocate distinct models for different cancer types regardless of which organ the tissues were collected from. Lastly, our dataset combination may be used to develop a universal model that not only classifies cancer types accurately but also provides genetic information as in the study by Fu *et al*^13^ who merged 28 TCGA cohorts to train a model for tissue/cancer classification that also provides genetic and survival information.

The current study has a few limitations. All patches were labeled corresponding to the slide labels resulting in noisy targets for supervised learning. This may have been problematic because some patches used to train the positive cohort discrimination model *D_p_* and cancer detection models *C_s_* may not have contained cancer despite the fact that they were labeled otherwise. We may have obtained different results had we used a weakly supervised algorithm or if we had acquired precise annotations by pathologists. The quality of patches was inconsistent; if we had sampled patches only near the center of the slide, with more slides available for training/testing to account for the deficient number of training/testing samples, we may have reached different conclusions.

In conclusion, we showed that deep learning algorithms can provide objective measures of morphological similarity among multiple cohorts, and that based on this similarity among cohorts, aggregating cohorts can be done successfully for more efficient development of high-performance diagnostic deep learning algorithms.

## Materials and Methods

### Dataset

TCGA slide images were used to train and test our cancer detection models. Despite the higher quality of formalin-fixed paraffin-embedded (FFPE) images, we used only the frozen tissue images since the TCGA dataset contains an insufficient number of negative FFPE images. Among the cohorts present in the TCGA dataset, cohorts with less than 36 positive (cancer present) and 36 negative (cancer absent) slides were removed, leaving a total of 12 types: KIRC, Kidney renal clear cell carcinoma; LIHC, Liver hepatocellular carcinoma; THCA, Thyroid carcinoma; OV, Ovarian serous cystadenocarcinoma; LUAD, Lung adenocarcinoma; LUSC, Lung squamous cell carcinoma; BLCA, Bladder Urothelial Carcinoma; UCEC, Uterine Corpus Endometrial Carcinoma; BRCA, Breast invasive carcinoma; PRAD, Prostate adenocarcinoma; COAD, Colon adenocarcinoma; STAD, Stomach adenocarcinoma (Table 1).

To balance the number of positive and negative samples for each cohort in training the cancer detection models, we sampled 5 sub-datasets as follows: each sub-dataset was sampled uniformly at random such that each sub-dataset contains 18 positive and 18 negative slides in each cohort for training and the same number of slides for validation/testing without overlapping slides for a total of 72 slides per cohort. The cohort discrimination models were also trained using a similar dataset partition: 18 and 36 slides per cohort for the positive/negative discriminative and general (positive and negative) models, respectively, and half for each validation and test sets. All experiments were conducted independently on the 5 sub-datasets to obtain statistically significant conclusions.

### Training / Inference

All models shared the same ResNet-50 V2 architecture operating on patches evenly cropped from slides with spatial resolution Ω = {1, …, 224}^2^, where each pixel spans 1.2*μ* meters, and were trained to make slide-level predictions for both cohort discrimination and cancer detection tasks^21^. For training, each patch’s label was assigned its corresponding slide-level groundtruth (i.e. cohort type for the cohort discrimination task and cancer presence/absence for cancer detection). Upon inference, the slide-level prediction 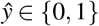 was computed using a likelihood-ratio test 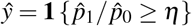 for some threshold *η* ≥ 0 that determines the operating point on the ROC curve, and the model’s pixel-wise predictions [*p_y_*(*i*, *j*)]_(*i,j*)∈Ω_ were summed channel-wise to obtain the class confidences 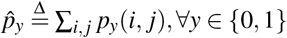.

All models were trained using the Adam optimizer with learning rate 10^−3^ until the validation accuracy saturated for 5 epochs with batch size of 32. Data augmentation was performed using the following: all input patches were rotated by multiples of 90 degrees and repeated to the horizontally flipped patch; pixel-level perturbations were performed randomly in the following order with maximum brightness change of 64/255, saturation ≤ 0.25, hue ≤ 0.04, contrast ≤ 0.75, and the resulting pixels were clipped to values in [0, 1]. Our implementation was based on the DL framework TensorFlow^22^.

### Cohort discrimination models and domain adaptation/generalization

The cohort discrimination tasks resemble the 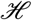-divergence 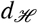 introduced for a remote task known as domain adaptation and generalization^14^. In particular, this metric in its original context quantifies the disparity between two domains (in our context, cohorts) characterized by their distributions *P_i_* and *P_j_* over possible images. Borrowing from its original context, Ben-David showed that a small 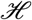-divergence between two cohorts characterized by *P_i_* and *P_j_* signifies that a model trained on an aggregate of cohort i and j will likely achieve small error when tested on either cohort. An exact computation of 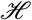-divergence requiring infinite validation samples is impossible, but its estimate 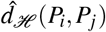 computed over a finite sample size can be used instead. The general discrimination model directly estimates the 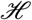-divergence in a pairwise manner, i.e. if instances from cohort *P_i_* are often (mis-)classified as coming from *P_j_*, cohort i is similar to cohort j. In contrast, the positive and negative discrimination models are conditional variants of 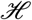-divergence, conditioned on the fact that the input image is either positive or negative.

### Aggregating Cohorts

Hierarchical clustering analysis (HCA) was performed using Ward’s method with the Euclidean distance between cohorts represented by their performances (rows) for each task, e.g. AUROC of single cancer detection model on target cohorts. The orderings were then used to displace cohorts in the similarity matrix such that neighboring cohorts indicate HCA clusters. After determining the dataset configuration, i.e. which cohorts to aggregate, its cohort grouping is reserved for a cancer detection model to be trained and tested on to obtain the AUROC performances as in Figure 4.

### Visualization Details

The uniform manifold approximation and projection (UMAP) visualization was attained using custom parameters (number of neighbors = 20, minimum distance = 0.5) on the features extracted from the penultimate layer of the universal model for sub-dataset 1. Interpreting the visualization is difficult when using excessive number of slides, and we instead randomly sample 20 patches per slide from the training sub-dataset to extract the features for this visualization.

## Author contributions statement

J.P., D.I.K. and K.H.J. conceived the experiment(s). J.P., S.T.K., Y.W.K. and K.K. conducted the experiment(s). S.T.K. helped design the experiments. J.P., Y.R.C., S.T.K. and H.P. analysed the results. J.P., Y.R.C. and S.T.K. wrote the manuscript. All authors reviewed the manuscript.

